# Network-based metabolite ratios for an improved functional characterization of genome-wide association study results

**DOI:** 10.1101/048512

**Authors:** Jan Krumsiek, Ferdinand Stückler, Karsten Suhre, Christian Gieger, Tim D. Spector, Nicole Soranzo, Gabi Kastenmüller, Fabian J. Theis

**Author notes:** co-first authors (Jan Krumsiek, Ferdinand Stückler).

## Abstract

Genome-wide association studies (GWAS) with metabolite ratios as quantitative traits have successfully deepened our understanding of the complex relationship between genetic variants and metabolic phenotypes. Usually all ratio combinations are selected for association tests. However, with more metabolites being detectable, the quadratic increase of the ratio number becomes challenging from a statistical, computational and interpretational point-of-view. Therefore methods which select biologically meaningful ratios are required.

We here present a network-based approach by selecting only closely connected metabolites in a given metabolic network. The feasibility of this approach was tested on *in silico* data derived from simulated reaction networks. Especially for small effect sizes, network-based metabolite ratios (NBRs) improved the metabolite-based prediction accuracy of genetically-influenced reactions compared to the ‘all ratios’ approach. Evaluating the NBR approach on published GWAS association results, we compared reported ‘all ratio’-SNP hits with results obtained by selecting only NBRs as candidates for association tests. Input networks for NBR selection were derived from public pathway databases or reconstructed from metabolomics data. NBR-candidates covered more than 80% of all significant ratio-SNP associations and we could replicate 7 out of 10 new associations predicted by the NBR approach.

In this study we evaluated a network-based approach to select biologically meaningful metabolite ratios as quantitative traits in GWAS. Taking metabolic network information into account facilitated the analysis and the biochemical interpretation of metabolite-gene association results. For upcoming studies, for instance with case-control design, large-scale metabolomics data and small sample numbers, the analysis of all possible metabolite ratios is not feasible due to the correction for multiple testing. Here our NBR approach increases the statistical power and lowers computational demands, allowing for a better understanding of the complex interplay between individual phenotypes, genetics and metabolic profiles.

## Introduction

Genome-wide associations studies (GWAS) provide valuable insights into genetic influences on complex diseases and phenotypes. Many risk loci were identified in recent studies [1], the mechanistic interpretation of their results, however, is often difficult [2]. Effects which cause a specific phenotype are often a mixture of many underlying processes. To overcome these limitations, GWAS with quantitative metabolic traits (mGWAS) were introduced to analyze genetic factors that have an effect on intermediate processes which are then referred to as genetically-influenced metabotypes (GIM) [3]. When applied for instance to selected metabolic pathways in plants [4,5] and large-scale data from human population cohorts [6-12], the mGWAS approach yielded new metabolite quantitative trait loci (mQTL), i.e. genetic regions which affect the levels of metabolites. Recent studies revealed direct associations between metabolic traits and genetic variants located near to genes encoding metabolite-specific enzymes or transporters [6,7]. For example, a single-nucleotide polymorphism (SNP) in the N-acetyltransferase 8 (*NAT8*) locus was reported to associate with N-acetylornithine [13].

Using metabolite concentration ratios as quantitative traits in addition to single metabolite concentrations further improved the results and interpretation of SNP-metabolite associations. It was shown that ratios between metabolite concentrations pairs reduced the overall biological variability in population data and resulted in robust statistical associations [5,14]. For instance, the level of a nutritional metabolite, but also the respective breakdown products, might be elevated in specific subjects. The metabolite ratio between these metabolites accounts for this inter-individual variation. In a biochemical interpretation, the ratio between product-substrate metabolite pairs can be interpreted as a proxy of the corresponding enzymatic reaction rate [15]. For example, Suhre *et al.* [13] reported that the association of a genetic variant in the *FADS1* locus and the ratio between fatty acids 20:3 and 20:4 is much stronger compared to the association with the respective single metabolite levels. The *FADS1* locus encodes for a fatty acid delta-5 desaturase with fatty acids 20:3 and 20:4 as substrate and product, respectively. The increase in association strength due to the ratio between reaction substrate-product pairs thus matches the biological function of the enzyme [16]. In previous studies all possible ratio combinations were tested for genetic associations. This “all-ratios” approach yielded associations that were of many orders of magnitude stronger compared to testing only individual concentrations [6,7,13].

However, taking all possible metabolite ratio combinations into account can be challenging from a statistical, computational and interpretational point-of-view, since inevitably many biochemically unrelated metabolite pairs are tested. In addition, the effect of a specific genetic variant on metabolites that are within a pathway might often be quite similar. Thus, conventional multiple testing approaches (like Bonferroni correction) might be too stringent for GWAS and possibly reduce the statistical power. While for large GWAS cohorts the limited power problem might be only an issue for small effect sizes, it can be beneficial for the design of case-control studies with small sample numbers. Additionally, the increasing number of measured variables will lead to a quadratic increase in the number of tests when dealing with ratios. Current methods allow for the detection of a few hundred metabolites, but this number will increase rapidly [17]. On a practical level it will be becoming computationally demanding to conduct association tests for up to 30 millions SNPs from several thousand individual genomes in multiple cohorts across millions of metabolite ratio combinations. The functional interpretation of the high-dimensional and complex data is also challenging for such study designs due to the vast amount of results produced. Preselecting meaningful ratio candidates based on biological network information thus is advantageous to overcome the above-mentioned limitations.

We asked whether it is beneficial to include prior information about the dependency between metabolites for the selection of biologically meaningful ratio candidates (network-based metabolite ratios, NBRs, see Fig. 1). Incorporating network information into the analysis of biological data has shown to be successful for example in proteomics and genomics applications [18-21]. On the level of a single pathway, we successfully applied the ratio approach for metabolomics data from a human challenging study, where we analyzed the beta-oxidation activity during a fasting period of 36 hours [22]. Here we could show that specific metabolite ratios which we derived from a model of fatty-acid breakdown revealed stronger associations to anthropometric and biochemical parameters such as BMI, body fat mass or insulin.

**Fig. 1.**
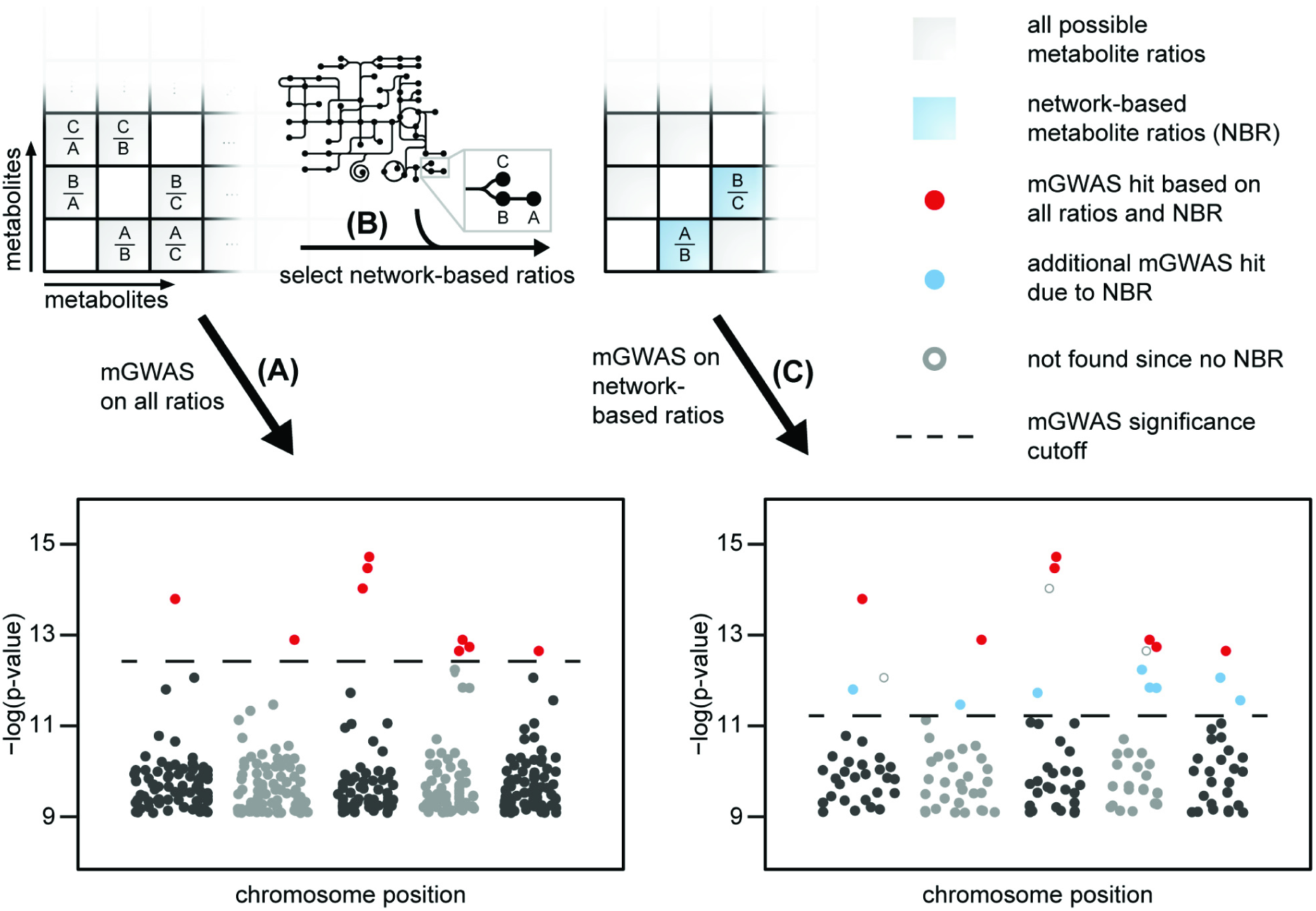
Network-based metabolite ratios (NBRs) for the analysis of genome-wide association studies with metabolic traits (mGWAS) Considering all possible metabolite ratios (red dots) as traits in mGWAS (A) has proven valuable in finding new functional insights about underlying biological processes [13]. Selecting network-based metabolite ratios instead of all possible ratios reduces the number of association tests and therefore results in a less stringent significance threshold after correction for multiple testing (B). Testing only for selected NBR (C) reveals new significant associations (blue dots). Due to incomplete metabolic network information or unknown complex regulatory effects some hits might not be found (grey circles).

For GWAS studies the metabolite dependencies need to be considered on a larger scale. This information can be obtained, for example, from metabolic pathways which are available from various sources such as KEGG, BiGG, EHMN and MetaCyc [23-26]. However, many measured metabolites are not annotated and the derived pathway information might be incomplete [27]. Statistical approaches like correlation-based methods, which purely rely on measured metabolomics data, can provide network information for all detected metabolites [28-30]. As metabolites in population data are highly correlated [31], we used partial correlations from Gaussian Graphical models (GGM) instead of normal correlations. Here indirect interactions between metabolites are removed before using the data-driven network for selecting ratio candidates.

Using metabolite concentrations or ratios as quantitative traits has generated novel hypotheses for biological functions of genetic loci. In the present article, we show how metabolic networks can be used to analyze the mGWAS data and facilitate the functional characterization of the respective results. The manuscript is organized as follows: First we test the NBR approach on simulated population data. We then evaluate whether ratio-SNP association hits reflect metabolic pathway reactions based on mGWAS results of a human population cohort. To this end, we test if metabolites that have a significant ratio-SNP association are more closely connected in metabolic networks reconstructed using GGM. In addition, we compare mGWAS results that were obtained using the ‘all ratio’ approach with our NBR method. For this comparison we first consider NBRs that were selected based on metabolic networks from databases. Since such derived networks are incomplete due to missing annotations, we also use networks reconstructed in a purely data-driven fashion from metabolomics measurements using Gaussian graphical modeling. Furthermore, we discuss newly predicted associations and their replication in an independent study cohort. In addition, we analyze associations which are not detected by the NBR approach in the context of pathway-related metabolites that are all affected by the same genetic locus.

## Results

### Network-based metabolite ratios improve mGWAS analysis of simulated reaction networks

Simulated reaction networks are useful tools to investigate the properties of biological systems and to examine new approaches in a well-defined setup [31,32]. We used such a framework to address whether selecting network-based metabolite ratios improves the SNP-ratio associations results, compared to taking all possible ratios. To this end, we computationally generated metabolomics measurements resembling features of a real population (see Fig. 2 A for a scheme of the simulation and methods for a detailed description of the model parameters).

**Fig. 2.**
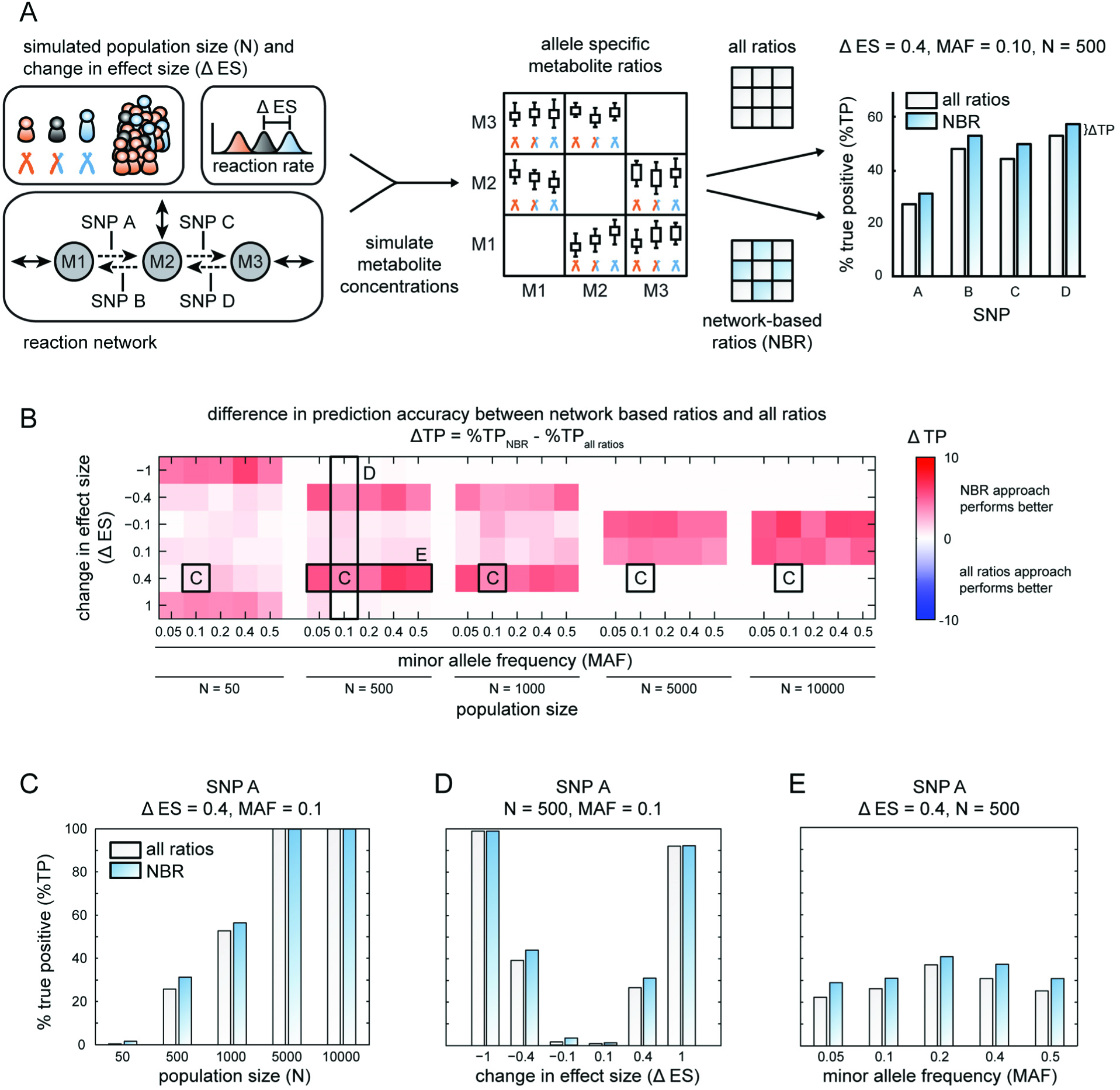
Network-based metabolite ratios (NBR) on simulated reaction networks. The NBR approach improves the analysis of ratio-SNP associations. A: Scheme of *in silico* simulation of SNP effects in metabolic reaction networks. For specific population sizes (N), minor allele frequencies (MAF), SNP effect sizes (ES) and reaction network topologies with different SNPs, steady state metabolite concentrations were simulated. Based on selected metabolite ratio sets (all ratios or PW-NBR) SNP-ratio associations were calculated. For true positive prediction, the best association hit matches the underlying reaction in the network. The fraction of truly predicted associations (%TP) was evaluated from 500 iterations. Since less association tests are needed using NBR, this approach is more sensitive, reflected by higher %TP values. B: Differences in %TP between PW-NBR and ‘all ratios’ analysis (ΔTP). The simulation was based on the reaction network depicted in A with one SNP-affected reaction between M1 and M2 (SNP A). A more detailed view of specific scenarios is given in subfigures C, D and E. Especially for small sample numbers in combination with small effect sizes the NBR approach improves the association analysis. C, D, E: Simulation results for selected scenarios as marked in B, with varying population size (C), effect size differences (D) or minor allele frequencies (E).

Our model incorporates genetic variation that has an effect on the respective enzyme activities. Such variation has been reported for example in a GWAS study that found an association between several SNPs in the *ACE* structural gene and ACE activity [33]. The reactions that we studied followed mass-action kinetic rate laws and were implemented as ordinary differential equations [31]. In order to account for variation between individuals, each reaction rate was drawn from a log-normal distribution [34] and then used to calculate individual steady state metabolite concentrations. Metabolites involved in SNP-affected reactions and their corresponding ratios showed genotype-specific levels with linear dependency, which is in accordance with previous studies on real data [6,7,13,35,36] (see S1 File for an example of simulated genotype-dependent metabolite ratios).

For all association tests, metabolite ratio candidates were selected by three different approaches: 1) all possible ratio combinations between all metabolites (‘all ratios’), 2) only ratios between connected metabolites in the network assuming that we know all true pathway reactions (network-based metabolite ratios from pathway information, PW-NBR) and 3) only ratios between neighboring metabolites in the network reconstructed from simulated metabolomics data using Gaussian Graphical modeling (network-based metabolites ratios from GGM information, GGM-NBR) [31]. Results for the evaluation of approach 2 can be found in Fig. 2; results for further network topologies are discussed in S2 File.

We tested for SNP-ratio associations using a linear model with genotype as independent variable and the respective metabolite ratio as response. All SNP-ratio associations were adjusted for multiple testing using Bonferroni correction based on the approach-specific number of ratio candidates. A ratio was counted as true positive if the best SNP-ratio association hit (lowest p-value) matched the underlying network, meaning that the simulated SNP was affecting the direct reaction between the two ratio metabolites. For instance, if the ratio M1/M2 in the metabolic network as depicted in S1 File shows the best association to SNP A, this SNP-ratio association is true positive, since SNP A directly affects the reaction between M1 and M2. As a quality measure the fraction of truly predicted associations (%TP) compared to all predictions was calculated.

Using this simulation framework, we tested several network topologies with different SNP-affected reactions and varying population sizes (N), minor allele frequencies (MAF) and SNP effect sizes (ΔES). The schematic workflow is depicted in Fig. 2 A for a linear reaction network consisting of three metabolites connected by reversible reactions. For each species we introduced exchange reactions, reflecting interactions with other metabolic pathways. The overview of all integrated scenario results in Fig. 2 B reveals that the network-based metabolite ratio approach performs equally well or even better compared to the ‘all ratios’ approach. NBR improves the prediction of SNP-reaction associations especially for scenarios with small effect sizes (ΔES = 0.4) in combinations with sample numbers of 500 and 1000. In order to detect small effects usually one has to increase the sample size, which is often a limiting factor. The improvement of the results is based on the different choice of ratio candidate sets. By only taking ratios of connected metabolites into account for the linear association model, we reduce the number of tests and increase the power of our analysis. Further examples for which the results of approach 1 and approach 2 are compared for different network topologies can be found in S2 File.

Fig. 2 shows the results for the NBR analysis with the given network structure from the simulated model (PW-NBR, see approach 2 above). As we are using a simulation framework, we know the underlying metabolic network and can easily determine neighboring metabolites for ratio selection. Since in reality most metabolic networks are not fully annotated and the PW-NBR approach may not be applicable, we also tested the third method (GGM-NBR) using reconstructed networks on the basis of the simulated steady state metabolite concentrations (S2 File). For linear cascades, PW-NBR and GGM-NBR show similar performance results. For more complex, branched reaction networks the prediction accuracy of PW-NBR is slightly better compared to GGM-NBR. Estimating the network based on the metabolomics data as done in the GGM-NBR approach here still performs better than the ‘all ratio’ approach. The simulation study shows that preselecting ratio candidates based on metabolic network information in most of the cases only improves the ratio-SNP predictions. These NBR improvements can especially facilitate the detection of small effects in studies with small sample numbers.

### Metabolite ratios significantly associated to specific SNPs are also closely connected in the metabolic network

We have shown on simulated metabolomics data that using network information about metabolite dependencies improves the analysis of genetically-influenced metabotypes. Next we tested the NBR approach on metabolomics and genotyping data based on 1,768 fasting serum samples from the German population study KORA [37] (“Kooperative Gesundheitsforschung in der Region Augsburg”), previously published in a genome-wide association study [13]. After quality control and stringent filtering the dataset contained measurements of 218 metabolites and 655658 genetic variants.

As discussed above, metabolite pairs whose ratio is significantly associated to a SNP should be closely connected in metabolic networks. In order to test this, we decided not to use networks based on pathways from databases due to missing or incomplete pathway annotations of many metabolites (122 out of the 218 metabolites could be mapped to KEGG, BIGG and EHMN [23-25]). Instead we used Gaussian Graphical modeling (GGM) to infer a pathway network for all 218 measured metabolites. Briefly, each edge in the network corresponds to a partial correlation coefficient above a certain threshold. Partial correlations represent pairwise correlations between metabolites after the confounding effects of all other metabolites and covariables have been removed. This approach has previously been shown to reconstruct pathways from blood serum metabolomics data in the same cohort [31,38]. Further information about the procedure, the metabolomics dataset and the obtained GGM can be found in [38].

Fig. 3 A shows the network representation of partial correlations in the GGM. Here metabolites which belong to a significant ratio-pair are marked red. We observe a clear grouping of pairs of red nodes in the network. For instance, the amino acids leucine, valine and glutamine are closely connected within the GGM, and are also part of ratios which are significantly associated to a SNP. We further asked whether metabolite pairs, which are both affected by the same genetic variant, are also closely connected in the metabolic network. To address this question, we compared the distribution of all pairwise metabolite shortest path lengths with the distribution of shortest path lengths between metabolite pairs whose ratio was significantly associated to a SNP (Fig. 3 B). To calculate the shortest paths, partial correlation coefficients were transformed to distance measures such that high partial correlation values then have low distances, meaning they are closely connected, and low partial correlation values are far apart (see Methods).

**Fig. 3.**
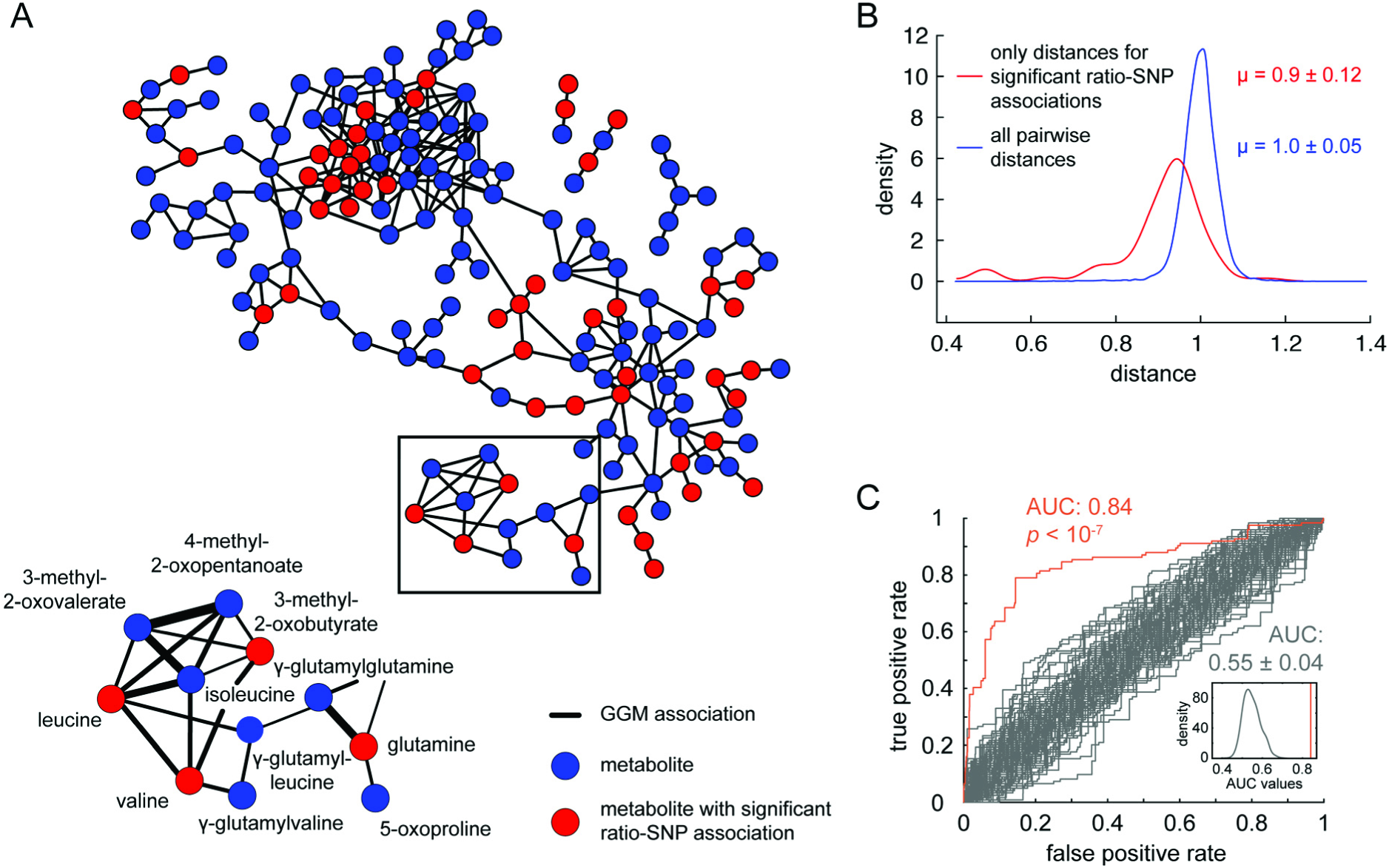
Analysis of GWAS with metabolite traits in the context of metabolic networks. Metabolite ratios that are significantly associated with specific SNPs are also closely connected in reconstructed metabolic networks. A: Network representation of a Gaussian Graphical model (GGM) reconstructed from large-scale metabolomics data as shown in [31,38]. Nodes represent metabolites and edges represent partial correlation values higher than 0.15. Zooming into the network reveals that the reconstruction puts metabolically related metabolites in a network context. Metabolites which belong to a ratio pair that is significantly associated to a SNP as reported in [13], are colored red. Line widths represent partial correlation strengths. B: Metabolite pairs, which are both affected by the same genetic variant, are also closely connected in the metabolic network. This can be seen using partial-correlation based shortest path distances between metabolites in the GGM. Compared to all distances in the GGM, significantly associated metabolite pairs tend to have smaller distances, i.e. higher partial correlation coefficients. C: ROC analysis of the distance separation seen in B. The area under the curve is 0.84 (orange line), compared to random networks (grey lines). The result is highly significant (empirical p-value < 10^−7^) and thus depends on the underlying GGM network used for the distance analysis.

Significantly associated metabolite pairs tend to have smaller shortest path distances, i.e. higher partial correlation coefficients, compared to all shortest path distances in the GGM. The mean distance between all metabolite pairs is 1, reflecting in our distance measure that most metabolites are not interconnected and have partial correlation coefficients close to 0. On the other hand, the average distance for metabolite pairs with significant ratio-SNP associations is 0.9 and thus more closely connected. ROC analysis [39] was used to quantify this separation, resulting in an AUC score of 0.84. In order to test whether this finding was only observed by chance or does indeed depend on the metabolite network structure, we compared our results to results obtained from randomized networks yielding AUC scores of 0.55 ± 0.04 (empirical p-value < 10^−7^). The ROC-analysis results for the original and randomized GGM network are shown in Fig. 3 C. This highly significant non-random AUC shows that most of the significantly associated metabolite pairs are in close distance. Our findings further suggest that we can use metabolic network information to preselect metabolite pairs for association studies of genetically-influenced metabotypes.

### Network-based metabolite ratios facilitate the analysis of mGWAS results by integrating genomic and metabolomics network information

The results from the toy simulation and the overall analysis of reported metabolite ratio-SNP associations demonstrated that network information can be used to select biologically meaningful ratio candidates for genetic association studies. In the following, we will address the question how to use this information in order to improve the analysis and interpretation of genetically-influenced metabotypes from mGWAS data. Since we want to focus our analysis on association hits at genetic locus level, we combined ratio-SNP associations that are within linkage disequilibrium of 0.8 or higher and only report the strongest hit (see Methods). Analogously to our simulation study, we used three approaches to select metabolite ratios for further association tests: ‘all ratios’, PW-NBR and GGM-NBR (see Fig. 4 A for a comparison of the results and S1 Table for a full list of all associations).

**Fig. 4.**
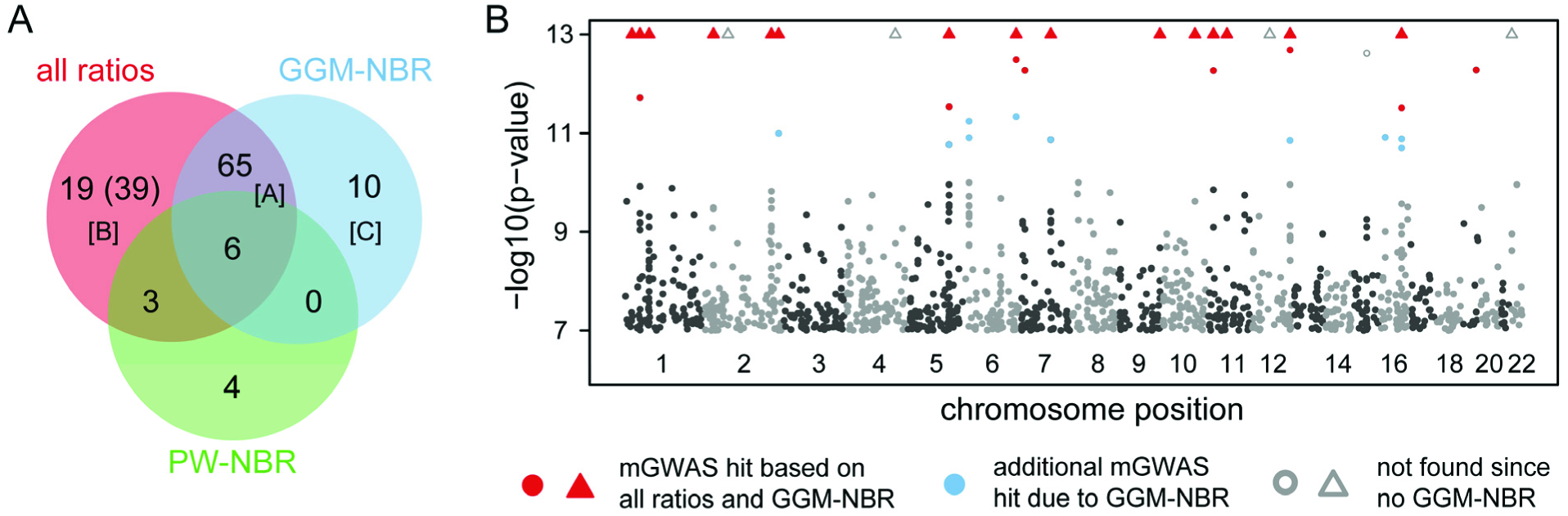
Comparison between results for different ratio candidate sets. A: Network-based ratio analysis yields similar associations compared to the ‘all ratios’ approach. Using reconstructed networks (GGM-NBR) performs much better compared to pathway-based networks (PW-NBR) due to the incomplete annotation of many metabolites. [A]: The overlap between the all-ratio and GGM-NBR approach associations is remarkably high. [B]: 20 out of 39 hits not identified by the network approach can be explained by pathway analysis of the underlying GGM network (see also Fig. 5 for two examples). [C]: Due to the reduced number of association tests and the resulting less stringent Bonferroni significance level, GGM-NBR reveals additional associations (see also Fig. 6). B: Manhattan plot of the results revealed by ‘all ratios’ and GGM-NBR approach. The strength of association for metabolite ratios is indicated as the negative logarithm of the p-value of the linear model. Only ratio-SNP associations with p-values below 10^−7^ are plotted. Triangles represent ratio-SNP associations with p-values below 10^−13^. Same ratio-SNP associations that are within linkage disequilibrium of 0.8 or higher are combined and only the strongest hit is shown. Significant mGWAS hits that were found by the ‘all ratios’ and GGM-NBR approach are marked in red (threshold after Bonferroni correction α = 3.22·10^−12^). Associations which are not detected by GGM-NBR are colored grey, while additional GGM-NBR results are marked as blue dots (threshold α = 2.01·10^−11^). Note that for our analysis we only considered reported associations between ratios and SNPs.

For the selection of meaningful network-based metabolite ratios we first used a pathway-based network (PW-NBR) that was built by combining information from KEGG, BIGG and EHMN [23-25]. Since not all metabolites are annotated in these databases, the network contains only 122 out of the 218 originally measured metabolites. Contrary to the *in silico* simulation study shown in Fig. 2, we do not have the full information about the true underlying reaction network to apply the PW-NBR approach for the mGWAS data set. We accounted for possible missing network connections by considering not only directly connected metabolites as ratio candidates, but also those with a network distance of one or two steps. The PW-NBR approach reveals only few significant SNP-ratio associations (13) and the overlap with the ‘all ratios’ approach is rather small (9 out of 113).

The GGM network on the other hand is purely data-driven and has the advantage of obtaining network dependencies for all 218 measured metabolites. The network was built by taking only metabolite pairs into account that showed an absolute partial correlation score of 0.1 or higher. We also tested other partial correlation cutoffs (S3 File), which however did not lead to a notable change of results. Similar to the PW-NBR analysis we also accounted for missing connections by selecting metabolites as ratio candidates that were connected in the network via one or two steps. The GGM-NBR approach reveals 81 significant SNP-ratio associations, which highly overlap with the results from the approach taking all possible ratios.

The comparison between ‘all ratios’, PW-NBR and GGM-NBR is shown in Fig. 4 A. The GGM-NBR approach yields considerably more significant ratio-SNP associations compared to the PW-NBR (81 vs. 13). This results from incomplete or missing annotations in pathway databases for almost 100 metabolites. In contrast, the full network information can be obtained for all measured metabolites using Gaussian Graphical modeling.

Though the overlap between ‘all ratios’ and GGM-NBR results is remarkably high (71 cases, set [A] in Fig. 4 A), there are some associations which are not observed using GGM-NBR (set [B], 39 cases). We inspected these cases in more detail by asking whether we could explain these effects by other effects of related metabolites. We hypothesized that in many cases the same underlying factor (e.g. genetic variation in one enzyme) influences metabolites in subsequent or neighboring reaction paths. To test this hypothesis, we calculated the shortest paths within the GGM network between the two metabolites of a not observed ratio-SNP association. On these paths we checked if there are other ratio pairs associated to SNPs in close genetic distance to the original ratio-SNP association. Two examples are shown in Fig. 5, one related to fatty acid metabolism and one related to sugar metabolism. Two SNP variants in the locus of *ACADM*, an enzyme of mitochondrial fatty acid beta-oxidation, for example are associated to different metabolite ratios (hexanoylcarnitine / acetylcarnitine and hexanoylcarnitine / oleate). On the basis of the data-driven reconstructed network, we can see that these metabolites are closely related. The effect of a genetic variant might affect specific metabolites, but also alter other metabolite concentrations within a pathway, both detected in mGWAS results. Using the GGM network context thus helps to understand these pathway effects, especially if no pathway dependencies for the metabolites of interest can be obtained from databases. In total we found 20 associations that could be explained by other associations within the same metabolic pathways, reducing the number of ‘all ratios’ only associations from 39 to 19. Hence by taking the network connections of metabolites into account we do not miss hits, but rather find the more direct associations, which point to the underlying biological mechanism.

**Fig. 5.**
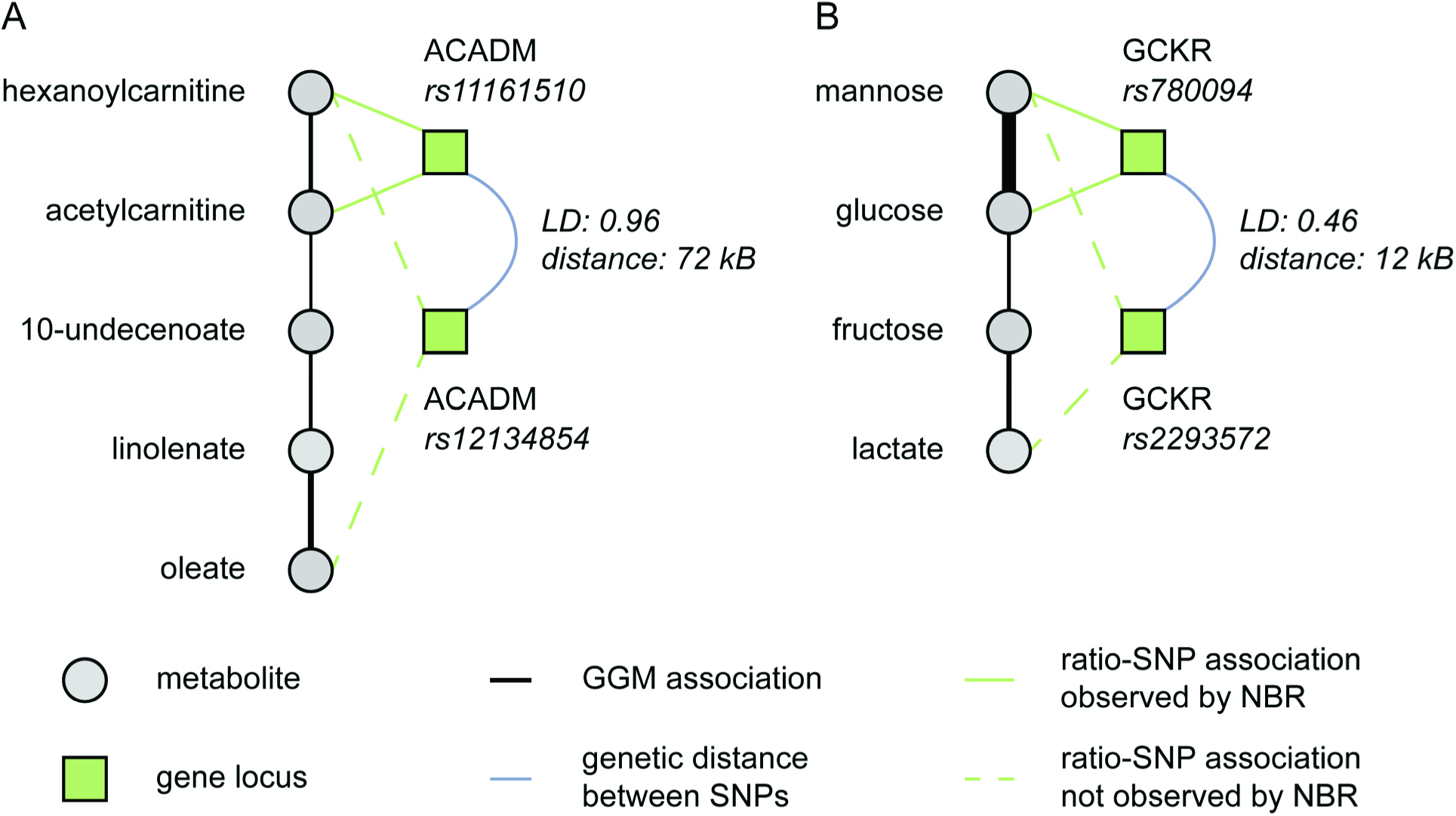
Metabolic network information reveals the interplay between different association loci. Analyzing associations within GGM-based metabolic paths helps to find the more direct associations which point to the underlying biological mechanism. Two examples are shown for fatty acid metabolism (A) and sugar metabolism (B). Genetic variants often affect related metabolite pairs. Two SNP variants in the ACADM locus for example are associated with different metabolite ratios. The respective metabolites are closely connected in the GGM network. Genetic effects within the ACADM locus thus have an impact on several metabolite concentrations within certain metabolic pathways. Line widths represent strength of partial correlation in GGM networks.

### Replication of associations predicted by the NBR approach

The GGM-NBR approach predicts 10 new ratio-SNP associations that were not found using the regular ‘all ratios’ approach (set [C] in Fig. 4 A and Manhattan plot in Fig. 4 B, see also Table 1). This results from a higher Bonferroni significance level (3.22·10^−12^ for ‘all ratios’ compared to 2.01·10^−11^ for GGM-NBR) due to fewer association tests. In S4 File we discuss the relationship between the number of measured metabolites and the number of ratios that have to be tested using the ‘all ratios’ and the NBR approach. 7 out of the 10 new ratio-SNP associations predicted by the GGM-NBR approach were replicated in the TwinsUK cohort data reported in [12], while none of the four associations predicted by PW-NBR could be replicated (Table 1 and S1 Table). Fig. 6 shows one example of a replicated genetically-influenced metabotype in the leucine metabolism. Using both the ‘all ratios’ and GGM-NBR approach, the ratio isovalerylcarnitine / isovalerate was found to be associated to the *OCTN2*/*SLC22A5* locus, which codes for an organic cation transporter with short-chain acyl esters of carnitine as substrates [40]. GGM-NBR analysis additionally revealed an association between isovalerylcarnitine / leucine and a SNP in the *ACSL6* locus. *ACSL6* catalyzes the formation of acyl-CoA species, possibly also isovaleryl-CoA, which is a degradation product of leucine but was not measured in the mGWAS study. While these associations might be based on indirect effects as isovalerylcarnitine is the transport form of isovaleryl-CoA, the network context in the GGM-NBR analysis helps to understand the interplay between different association loci and metabolite ratios. Table 1 displays further associations that were only found using GGM-NBR

**Fig. 6.**
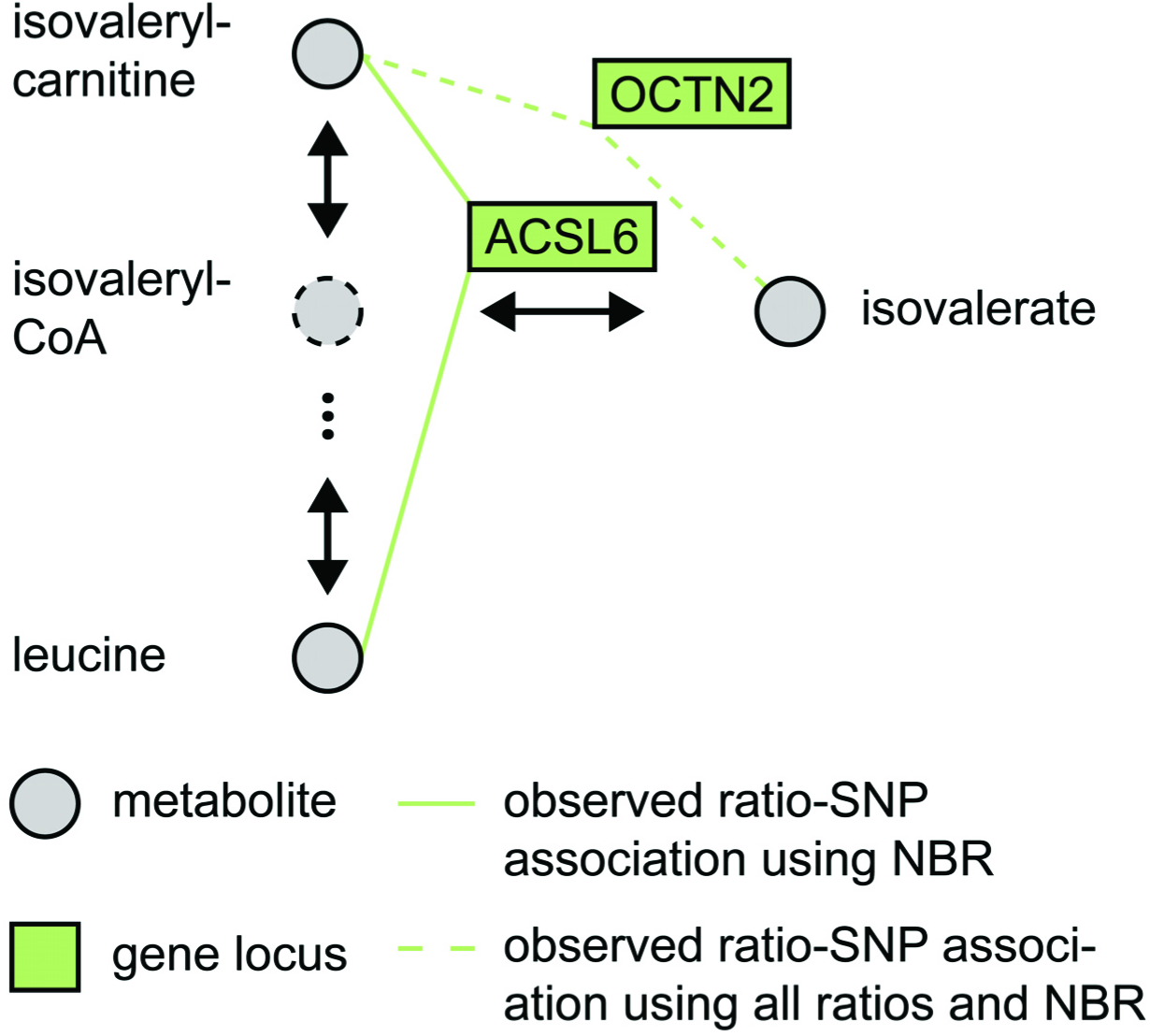
Example of an additional association found by the NBR approach. Both the ‘all ratios’ and NBR approach find an association between isovalerylcarnitine / isovalerate and the *OCTN2*/*SLC22A5* locus (rs274570), which codes for an organic cation transporter. Additionally, NBR-GGM analysis revealed an association between isovalerylcarnitine / leucine and a SNP in the *ACSL6* locus (rs10040809). *ACSL6* catalyzes the formation of acyl-CoA species like isovaleryl-CoA, which is a degradation product of leucine but was not measured in the mGWAS study (dashed circle). Metabolite relationships, which are obtained from known biochemical pathways or GGM networks, allow for a better understanding and interpretation of indirect effects and observed ratio-SNP associations. See Table 1 for a full list of all additional associations found by the NBR approach.

**Table 1:** List of all additional SNP-ratio associations found using the GGM-NBR approach. The associations between SNP rs9393903 and DHA/EPA ratio, as well between rs7200543 and 1-eicosatrienoylglycerophosphocholine/1-linoleoylglycerophosphocholine have been reported previously. However, this result was only obtained after increasing the sample size by combining two cohorts in a meta-analysis of two GWAS studies. Beta and p-value were derived from published association results of the KORA cohort [13] using a linear regression model based on additive genetic effects (using log10-scaled metabolic traits, genotype is coded as 0-1-2, major-hetero-minor genotype). X: Association was replicated in the TwinsUK cohort [12]. *: The two reported SNPs are both in the *ELOVL2* locus, but with linkage disequilibrium smaller than 0.8 (0.398) and therefore not combined. See also Fig. 6 for an example of a newly predicted association.

| Locus (SNP id) | Metabolite ratio | p-value | beta | Functional interpretation | Replication |
| --- | --- | --- | --- | --- | --- |
| *ACSL6*<br>(rs10040809) | isovalerylcarnitine / leucine | $1.70 \cdot 10^{-11}$ | -0.033 | ACSL6 catalyzes the formation of acyl-CoA species; isovaleryl-CoA is involved in leucine metabolism (see also Fig. 6) | X |
| *COX6A1*<br>(rs2076022) | butyrylcarnitine / propionylcarnitine | $1.40 \cdot 10^{-11}$ | -0.046 | COX6A1 is a terminal oxidase in mitochondrial electron transport; fatty acids are transported into mitochondria as acylcarnitines; ratio also associated to ACADS locus (observed by both all ratio and GGM-NBR) | X |
| *ELOVL2\**<br>(rs9393903 and rs3734398) | docosahexaenoate (DHA; 22:6n3) / eicosapentaenoate (EPA; 20:5n3) | $1.23 \cdot 10^{-11}$<br>and<br>$5.76 \cdot 10^{-12}$ | -0.030<br>and<br>-0.026 | EPA is substrate of ELOVL2, DHA biochemically related by desaturase reactions | X<br>and<br>X |
| *ZNF655*<br>(rs1581492) | androsterone sulfate / dehydroisoandrosterone sulfate | $1.35 \cdot 10^{-11}$ | -0.119 | gene encodes for a zinc finger protein; potential link to regulatory elements; ratio also associated to AKR1C isoforms (involved in androgen metabolism) | X |
| *HEATR7B1*<br>(rs10203853) | bilirubin (E;E) / oleoylcarnitine | $1.01 \cdot 10^{-11}$ | -0.047 | potential link to regulatory elements; ratio also associated to UGT1A, which has bilirubin as a substrate | X |
| *SLC7A6*<br>(rs6499172) | acetylcarnitine / glutaroyl carnitine | $1.30 \cdot 10^{-11}$ | -0.040 | SLC7A6 is involved in the transport of amino acids and also associated to glutaroyl carnitine/ lysine ratio; possible pathway interaction | - |
| *PRMT7*<br>(rs2863978) | acetylcarnitine / glutaroyl carnitine | $1.99 \cdot 10^{-11}$ | -0.034 | PRMT7 is a arginine methyltransferase for protein modification; potential link to regulatory elements | - |
| *SLC22A1*<br>(rs456598) | gamma-glutamylvaline / isobutyrylcarnitine | $4.65 \cdot 10^{-12}$ | 0.064 | SLC22A1 is a transporter for many organic cations; SNP associated to serum concentrations of total cholesterol and low-density lipoprotein cholesterol [41] | - |
| *PDXDC1*<br>(rs7200543) | 1-eicosatrienoylglycerophosphocholine / 1-linoleoylglycerophosphocholine | $1.22 \cdot 10^{-11}$ | -0.035 | Association suggests that PDXDC1 is involved in the glycerophosphocholine-metabolism | X |

## Discussion

Genome-wide association studies with metabolite ratios as quantitative traits have deepened our understanding of the complex relationship between genetic variants and observed phenotypes. It has been shown that ratios between metabolite concentrations pairs reduce the overall biological variability in population data resulting in robust statistical mQTL associations [14]. In previous studies, metabolite ratios were either manually selected with respect to specific enzymatic reactions [8] or all possible ratio combinations were used [6,7,13]. Especially due to the large number of all possible ratios for studies with many metabolites it is important to narrow down the number of association tests. We argue that the proper selection of ratio candidates based on metabolic network information will improve the analysis of association studies with metabolic traits from a statistical, computational and interpretational point-of-view.

In this work we propose to choose biologically meaningful metabolite ratios based on metabolic networks. In a study on the dynamics of human metabolism, we applied this concept on the level of a single pathway and used a fatty acid beta-oxidation model to infer metabolite ratios reflecting enzymatic activity [22]. For the analysis of mGWAS results, we here extended this approach by going from local pathways to global metabolic interaction networks to select network-based ratios (NBRs) for further association tests.

Before applying NBRs on human population GWAS data, we used simulated reaction networks. Our model simulates differences in metabolite levels, which result from genetic variation affecting enzyme activities. Such effects have been reported for several SNPs in the *ACE* structural gene and the ACE activity [33]. It is to be noted that the *in silico* model is obviously an oversimplified model of gene-metabolite interactions. The primary goal of the presented analysis was to test our hypothesis in a well-defined and comprehensible environment before going to noisy experimental data. Our *in silico* results show that the NBR approach is applicable for small sample size studies and, even more important for practical applications, for genetic variants with small effect sizes.

We further analyzed mGWAS results from two different study cohorts as an additional evaluation of the NBR approach. Initially the mGWAS results were obtained by using all possible metabolite ratio combinations as traits. We compared the associations detected using all ratios with associations observed after testing only ratio candidates derived from metabolic networks. Data-driven metabolic networks (GGM-NBR) gave similar results as the ‘all ratios’ approach. 7 out of 10 new associations predicted by the GGM-NBR approach could be replicated in a much larger study, suggesting that information about metabolite dependencies from data-driven networks allows for detecting associations in studies with smaller sample size. Networks obtained from pathway annotations in the literature (PW-NBR) could not reveal many associations. The limited results from PW-NBR are certainly based on the sparseness of metabolite annotations and network information in databases like KEGG, BiGG and EHMN. Data-driven reconstruction methods provide the opportunity to measure relations between all measured metabolites. It is important to acknowledge that results from statistical inference methods like Gaussian Graphical models should not be confused with a perfect reconstruction of metabolic pathways [42]. For instance, if intermediate metabolites cannot be detected, connections in reconstructed networks do not necessarily represent direct biochemical pathway reactions. With advanced metabolite detection techniques and network reconstruction methods, both the annotation and the data-driven pathway information will further improve and the two network sources can be combined to enhance our understanding of metabolism by finding more genetically-influenced metabotypes. For new datasets, ratio candidates then can be selected both from established candidate sets of previous studies, as well from de-novo calculated networks based on the new data.

Narrowing down the size of ratio candidate sets is also important for small, phenotype-specific studies. For example, studies investigating rare variants or small effect sizes often have to deal with small case numbers. For such studies it is essential to reduce the number of tests in order to improve the statistical power and lower computational demands. Moreover, advanced metabolomics methods will soon allow for the detection of several thousand metabolites. At this point it will not be feasible anymore to test all possible ratio combinations against genetic variants in order to find genetically-influenced metabotypes. Testing only for selected ratios circumvents these dimensionality problems, since the number of network-based ratios does not increase so rapidly (see S4 File). In addition, we could show that genetic effects in one locus have an impact on the concentration levels of biochemically related metabolites. It is important to understand at this point that measuring the distance between metabolites is not straightforward, both in reconstructed and literature-based metabolic networks [43,44]. Nevertheless, the network context helps to find the more direct associations which point to the underlying biological mechanism.

The NBR approach may also be combined with methods accounting for the inherent correlation between SNPs due to linkage disequilibrium [45,46], thus reducing the number of both ratios and SNPs for multiple testing correction. The presented approach is not restricted to association studies with metabolic traits and can be extended to other quantitative *omics* data, also in case-control studies. For such studies the sample size is usually much smaller and our preselection of ratios could improve statistical power. Moreover, NBRs can be used for other quantitative biomolecular data such as gene expression measurements or epigenetic modifications [47]. Here the interactions between gene products might be both inferred from data or obtained from biological pathways, well-established protein-protein or gene-regulatory networks [20,21,48].

## Conclusions

GWAS with metabolite ratios as quantitative traits have deepened our understanding of genetic effects on metabolic functions. Usually all metabolite ratio combinations are tested for associations to genetic variants, which can be challenging from a statistical, computational and interpretational point-of-view. For data with many metabolites, taking metabolic network information into account is of great benefit for the analysis and interpretation of association results. Our network-based metabolite ratio approach reveals nearly the same associations compared to the ‘all ratios’ approach and we could replicate 7 out of 10 new associations predicted by the NBR approach. Using NBR allows for the detection of weaker effects, since considering only biologically meaningful ratio candidates increases the statistical power. For upcoming studies with large-scale metabolomics data and small sample numbers, our NBR approach provides a valuable tool to increase the statistical power, lower computational demands and facilitate the interpretation of the results. Network-based analysis will then help to better understand the complex interplay between individual phenotypes, genetics and metabolic profiles.

## Methods

### In silico simulation of SNP effects on metabolic reaction networks

Metabolic reaction networks were simulated using mass-action kinetics. A dynamical system with *m* metabolites and *r* reactions can be represented using the stoichiometric matrix *S^m×r^*, where each row represents a compound and each column a reaction. Negative entries in *S* represent educts, positive products of one specific reaction. The change of one metabolite over time is then described by a system of linear equations

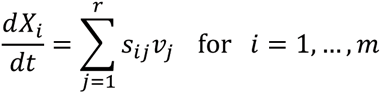

with rate *v*_*j*_ for the *j*th reaction and (s_ij_) = *S* for the entries of the stoichiometric matrix. The topology of the reaction network thus is fully represented by *S*, while the dynamical properties are determined by the reaction rates *v* according to the law of mass action kinetics [31]. Steady state metabolite concentrations are obtained either by solving the system of linear equations (i.e. setting all equations to zero) for linear systems or by simulating the dynamical system until it has reached its equilibrium.

Using this framework, different scenarios for reaction networks topologies, altered reaction rates due to SNP effects, allele frequencies and population size were investigated. The allele frequencies were calculated following Hardy-Weinberg equilibrium model with chosen minor allele frequency between 0.05 and 0.5. To account for variability in the simulated population, the reaction rates for each individual were randomly and independently drawn from log-normal distributions with mean 3 and standard deviation 1. The effect of a specific SNP was modeled by adding the effect size once (heterozygote case) or twice (homozygote minor allele case) to the mean log-normal parameter of the affected reaction rate. For example ΔES = 0.4 indicates that the rates for the major allele homozygote, heterozygote and minor allele homozygote case are drawn from log-normal distributions with mean 3.0, 3.4 and 3.8, respectively. The population size was chosen to be between 50 and 10,000. We used a multiplicative error model in order to take technical noise into account, which arises during the measurement of metabolomics samples. To this end, calculated metabolite steady state concentrations were multiplied with random factors drawn from a log-normal distribution with mean 1 and standard deviation 0.05. As a result we obtained all metabolite concentrations for each individual in the population depending on its genetic background.

### Reconstruction of metabolic networks using Gaussian Graphical modeling

Gaussian graphical models (GGM) are calculated using full-order partial correlation coefficients, i.e. each pairwise correlation is corrected against all remaining (n-2) variables to remove indirect effects. For data with more samples than variables, full-order partial correlations can be calculated by a matrix inversion operation. A more detailed description of GGM calculation based on metabolomics data can be found in [31]. Since in our simulated data there are for some cases less samples than variables (metabolites), we used the Rpackage GeneNet [49] which calculates a regularized version of partial correlation coefficients. This method yields also for cases with more samples than variables robust estimates of partial correlation coefficients. All computations were performed on log-transformed metabolite concentrations, as testing for normality revealed that for most cases the log-transformed concentrations were closer to a normal distribution than the untransformed values [13].

### Association between metabolite ratios and SNP effects for simulated reactions networks

The simulated metabolite concentrations showed log-normal distributions and were therefore log-transformed for further statistical analysis. For all association tests, metabolite ratio candidates were selected by three methods: 1) all possible ratio combinations between each metabolite in the underlying reaction network, 2) only ratios between neighboring metabolites in the reaction network and 3) only ratios between neighboring metabolites in the network structure reconstructed from metabolomics data using GGMs. The third approach represents a purely data-driven approach that does not require known pathway interactions as input and is thus independent of functional annotations of the measured molecules (see description above). We used a linear regression model based on additive genetic effects to test for the association between metabolite ratios and genetic background in the simulated concentrations, as previously reported in several GWAS studies [6,13,35], with genotype as independent variable and the respective metabolite ratio as response. Regression coefficients were tested for significant deviation from zero. To account for the number of tests for each ratio candidate set, Bonferroni correction was applied. If the best SNP-ratio association (lowest p-value) matched the underlying reaction networks, the test was counted as true positive. In this case the simulated SNP was affecting the direct reaction between the two ratio metabolites. The fraction of truly predicted associations (%TP, number true positive cases divided by all cases) was used to assess the quality of each ratio candidate selection method.

### Analysis of metabolic distances in Gaussian Graphical models

The distance d_i,j_ between metabolites M_i_ and M_j_ in the GGM was calculated based on the respective partial correlation coefficient ζ_i,j_, which was transformed by d_i,j_ = exp(−ζ_i,j_). Closely connected metabolites with high partial correlation coefficients have small distances. Based on this distance measure we calculated shortest paths between all metabolite pairs. We compared the distribution of all pairwise metabolite shortest path lengths with the distribution of shortest path lengths between metabolite pairs whose ratio was significantly associated to at least one SNP (adjusted Bonferroni threshold *p*<3.2241.10^−12^). ROC analysis [39] was used to quantify the separation of the two distributions. To assess the significance of this observed AUC score, we performed graph randomization by edge rewiring on the distance-weighted graph as described in [50]. During each randomization step the target nodes of two randomly chosen edges are exchanged. In order to achieve sufficient graph randomization, the exchange step is repeated five times the number of edges in the graph, as suggested in [51]. For the empirical p-value calculation we performed the distance-based ROC analysis for 10^7^ randomized graphs.

### NBR for GWAS data

We established our network-based ratio approach on data from the KORA population cohort. Details about the sample acquisition, metabolomics measurements and genotyping can be found in [13]. Briefly, we used metabolomics measurements of 295 metabolites and genotyping data for 655,658 SNPs from 1,768 fasting serum samples of the KORA study. Quality control of metabolomics data and stringent filtering resulted in 218 metabolites that were used for further analysis. Since all metabolomics data were transformed to log-scale for further statistical tests, we calculated metabolite ratios by taking the difference between the log-transformed concentrations, yielding 23,653 metabolite ratios for the ‘all ratios’ case. Since we want to focus our analysis on association hits at genetic locus level, we combined ratio-SNP associations that were within linkage disequilibrium of r^2^=0.8 or higher, based on LD data from HapMap derived from the SNAP server [52]. For cases where several SNPs within one locus were associated to the same metabolite ratio we only used the most significant association. No evidence of population stratification could be found in the population cohorts. Lambda values ranged from 0.965 to 1.024 (median 1.006) in KORA [6,13]. All participants in both TwinsUK and KORA have given written informed consent, and local ethics committees, the Guy’s and St. Thomas’ Hospital Ethics Committee for TwinsUK and Bayerische Landesärztekammer for KORA, approved the studies.

For the selection of ratio candidates based on network information we used two metabolic network sources: a pathway-based network (PW-NBR) and a GGM-based network (GGM-NBR). The first network was constructed by combining metabolite reaction information from three independent databases: 1) *H. sapiens* Recon 1 from the BiGG databases (confidence score of at least 4) [24], 2) the Edinburgh Human Metabolic Network reconstruction [25] and 3) the KEGG PATHWAY database [23]. Due to missing annotations, only 122 out of 218 measured metabolites were found in the combined pathway-based network. The GGM-based network is based on the network reported in [38] and was built by taking only metabolite pairs into account that showed an absolute partial correlation score of 0.1 or higher. In order to account for missing metabolic connections in the networks, we chose metabolites that were connected via one or two steps as ratio candidates for GGM-NBR and PW-NBR, resulting in 3,786 and 879 metabolite ratios, respectively.

As described above, a linear regression model based on additive genetic effects was used to test for the association between metabolite ratios and genetic background. The model was adjusted for age and gender as covariates. We applied Bonferroni correction to account for the large number of association tests. The p-value threshold was calculated by 0.05/(number of selected ratios * number of SNPs). Thus the adjusted threshold for genome-wide significance for the ‘all ratios’, GGM-NBR and PW-NBR analysis was *p*<3.2241.10^−12^, *p*<2.0142.10^−11^ and *p*<8.6757.10^−11^, respectively.

For SNP-ratio associations that were not discovered using the GGM-NBR approach we checked whether these effects could be explained by related metabolites. Based on the edge weights of the underlying GGM network we calculated shortest paths between the two metabolites of the ratio pair [53]. On these paths we checked if there are other ratio pairs which are associated to any SNP in close genetic distance to the original SNP, that was not found using GGM-NBR.

Replication of associations predicted by the NBR approach was performed using the data from the TwinsUK cohort results reported in [12]. In this study, we performed a genome-wide association analysis of metabolite ratio-SNP associations for 6,065 adult individuals similar to the analysis of Suhre *et al*. [13] in the KORA cohort which we used for the evaluation of the NBR approach (see above). The metabolomics data used for the replication has been measured on the same metabolomics platform as the data that we used for predicting new associations with our NBR approach. The replication was performed as follows: For each ratio-SNP association found in the KORA cohort we checked in the TwinsUK results for significant (p<2.0142·10^−11^, same cutoff as for GGM-NBR) associations between the ratio and SNPs with LD r^2^=0.8 or higher. For cases where several SNPs within one locus were associated to the same metabolite ratio, we only report the most significant association. As the replication was only performed for associations predicted by the NBR approach, it is to be noted that for a specific locus the here presented replicated association may not be the strongest association for this locus as reported in [12].

## Acknowledgements

The authors thank So-Youn Shin for assistance with the TwinsUK replication data. We thank all study participants of the KORA and the TwinsUK studies for donating their blood and time and all members of the field staff who planned and conducted the study.

## Supplementary Materials

- **Supplementary Material 1 - Genotype-specific metabolic traits in simulated reaction networks**
- **Supplementary Material 2 - Additional results for simulated reaction networks of different topologies**
- **Supplementary Material 3 - Evaluation of different GGM-NBR parameter settings**
- **Supplementary Material 4 - Estimating the number of ratio candidates for the ‘all ratios’ and GGM-NBR approach**
- **Supplementary Material 5 - List of all associations found in the KORA data and replication results based on TwinsUK data.**

